# Mechanisms of spectrotemporal modulation detection for normal- and hearing-impaired listeners

**DOI:** 10.1101/2020.01.03.894667

**Authors:** Emmanuel Ponsot, Léo Varnet, Nicolas Wallaert, Elza Daoud, Shihab A. Shamma, Christian Lorenzi, Peter Neri

## Abstract

Spectrotemporal modulations (STMs) offer a unified framework to probe suprathreshold auditory processing. Here, we introduce a novel methodological framework based on psychophysical reverse-correlation deployed in the modulation space to characterize how STMs are detected by the auditory system and how cochlear hearing loss impacts this processing. Our results show that young normal-hearing (NH) and older hearing-impaired (HI) individuals rely on a comparable non-linear processing architecture involving non-directional band-pass modulation filtering. We demonstrate that a temporal-modulation filter-bank model can capture the strategy of the NH group and that a broader tuning of cochlear filters is sufficient to explain the overall shift toward temporal modulations of the HI group. Yet, idiosyncratic behaviors exposed within each group highlight the contribution and the need to consider additional mechanisms. This integrated experimental-computational approach offers a principled way to assess supra-threshold auditory processing distortions of each individual.

## Introduction

An unresolved problem in auditory sciences concerns the large heterogeneity found among individuals at performing challenging auditory tasks, e.g. for understanding speech in the so-called “cocktail party” problem. Many factors ranging from sensory to cognitive likely contribute to this variability. This heterogeneity is typically found for people clinically diagnosed with sensorineural hearing loss (Moore, 2007). Surprisingly, recent studies have shown that this heterogeneity extends to middle-aged individuals with clinically-normal audiometric thresholds and similar cognitive resources (e.g. Ruggles et al., 2011). Such results have led to the view that suprathreshold auditory distortions not accounted for by pure-tone audiometry may have a substantial impact on speech-in-noise (SIN) understanding. Yet these distortions remain poorly characterized, despite their critical importance for understanding the SIN deficits typically found for these different groups of patients and for designing more effective hearing devices (Moore, 2007; Lesica, 2018).

Speech sounds convey salient spectral and temporal modulations (Singh & Theunissen, 2003; Varnet et al., 2017). Historically, psychoacoustical studies of suprathreshold auditory mechanisms have investigated these two dimensions separately (Miller et al., 2018) using signals modulated in instantaneous amplitude (AM), frequency (FM), or spectral ripples (Houtgast, 1989; Bacon & Grantham, 1989; Moore & Sek, 1996; Wallaert et al., 2018; Eddins & Bero, 2007; Ozmera et al., 2018). Results from these studies using AM and FM signals have led to the development of functional auditory models such as the (temporal) modulation filter-bank (Dau et al., 1997), the current gold standard for simulating suprathreshold processing in the auditory system (Biberger & Ewert, 2016). However, it remains unclear whether this model still constitutes a good account of how the auditory system processes complex signals showing *joint* spectral and temporal modulations as in the case of speech (Singh & Theunissen, 2003; Chi et al., 1999; Elhilali et al., 2003; Venezia et al., 2016, 2019); see Figure 1A.

**Figure 1.**
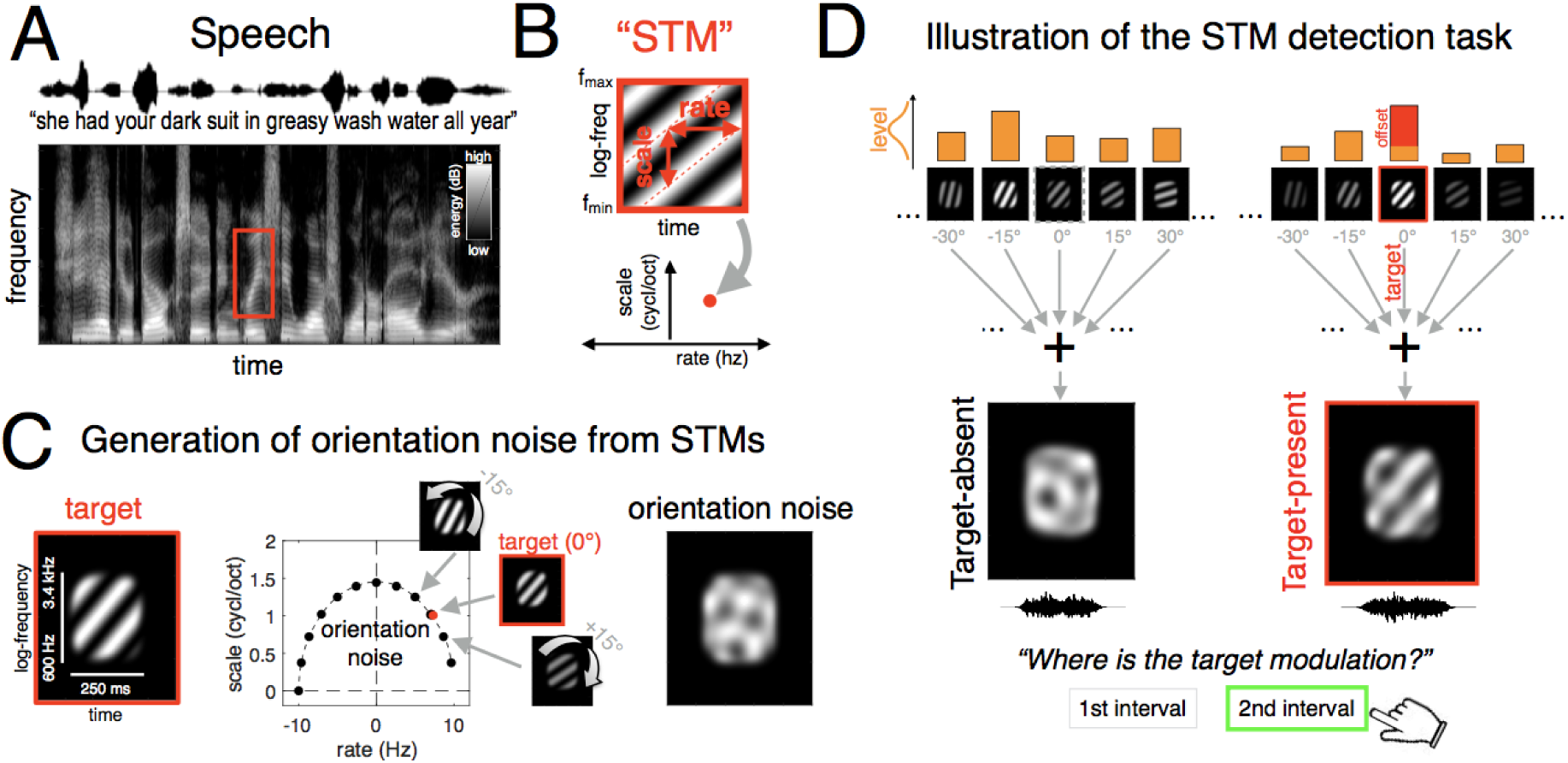
A) Example spectrogram of a speech sentence. Speech formants display clear spectrotemporal energy patterns (red box). B) Spectrotemporal modulations (STMs) or “ripples” are envelope modulations (top) specified by their spectral modulation density (cycles/octave, y axis) and their temporal modulation rate (Hz, x axis); they therefore correspond to a single point in scale-rate space (bottom). STMs represent good approximations to speech formants. C) Procedure for generating 1D orientation noise. The target stimulus (left) is an elementary STM (rate 7 Hz; scale 1 cycl/oct) oriented along the upward direction (from bottom-left to top-right). Orientation “ripple” noise is generated by summing 12 different ripple components made from 15° rotations of the target ripple in spectrotemporal space, each assigned random amplitude and phase. Each rotation corresponds to a different pairing of rate (from −10 to 10 Hz) and scale values (from 0 to 1.4 cycl/oct); this procedure specifies orientation noise (see example in right panel) as a 12-component vector spanning a 1D trajectory across rate/scale space (middle panel). D) Illustration of the STM detection task. The 12 levels specifying each noise sample of the target-absent stimulus (left) are drawn from a Gaussian distribution (only 5 are shown). The target-present stimulus is generated using the same procedure, except a constant level offset is added to the component corresponding to the target orientation (right panel, red offset). On each trial, listeners are presented with both stimuli in random order, and must determine which interval contained the target-present stimulus (bottom). Here the procedure is illustrated for an upward target, but we also tested (in different observers) detection of a downward target.

Spectro-temporal modulations (STM), also called “ripples”, may offer another description of critical speech features as they correspond to spectral modulations moving across the frequency scale over time (see Fig. 1B), thus constituting a first-order model of the spectrotemporal energy patterns of speech formants. It has been demonstrated that speech sounds can be well reconstructed and speech perception can be well accounted for using such signals (Chi et al., 1999; Elhilali et al., 2003; Mesgarani et al., 2006; Elliot & Theunissen, 2009). Physiological studies have provided converging evidence that the central auditory system is specifically tuned to process STMs (Hullet et al., 2016; Santoro et al., 2017) and the direct behavior relevance of STMs with rather low rates (1-10 Hz) and scales (1-2 cycles/octaves) to speech intelligibility has been demonstrated (Elliot & Theunissen, 2009; Venezia et al., 2016). Moreover, results from modeling studies indicate that cortical auditory models or metrics such as STMI (STM index), all based on a decomposition of auditory signals through a STM filter-bank, can accurately predict SIN intelligibility scores (Chi et al., 1999; Elhilali et al., 2003; Bernstein et al., 2013b). STMs may therefore constitute a unified approach for probing suprathreshold auditory processes as they are actually recruited by natural speech signals, and corresponding to how auditory information is finally represented and discriminated at a cortical level.

Recently, Oetjen and Verhey (2015, 2017) proposed a psychophysical masking paradigm to assess frequency selectivity to STMs at the behavioral level. All tested individuals were NH listeners, whose task was to detect a given STM target embedded within noise containing other STMs. By varying rate and scale of these masking modulations, they derived 2-D tuning functions in rate-scale space. Their results show that NH listeners do rely on bandpass filters in the STM domain, but that these filters are only partially able to differentiate upward from downward drifting modulations.

Because the above experiments involved NH listeners, we do not know how cochlear hearing loss impacts the characteristics of this process (tuning, directionality and possibly other characteristics). Most importantly, it is unknown how the auditory system makes use of different STM filters; the computational mechanisms that underpin the processing of spectral-only and temporal-only modulations have been explored in various studies (Dau et al. 1997; Ewert et al., 2002; Joosten et al., 2016), but the psychophysical mechanisms for joint spectral *and* temporal modulations have not been examined directly.

Several studies have started to address how STM signals are detected by both normal and impaired auditory systems (Bernstein, 2013a, 2013b, 2016, 2018; Merhaei et al., 2014; Miller et al., 2018). Importantly, they have shown that STM sensitivity can account for a significant proportion of the variance in speech-reception thresholds in noise (SRTs) for HI listeners, beyond that accounted for by the audiogram. Because this correlation was found with STMs at low modulation rates and high scales (Bernstein et al., 2016; Miller et al., 2018), these results point towards concurrent impairment of frequency selectivity (high scale) and processing of temporal fine structure (TFS) information in the stimulus waveform (low rate; Miller et al., 2018). However, they leave important questions unanswered: for example, they do not establish whether the observed deficits are primarily attributable to STM filtering anomalies, or whether signals are filtered normally but further represented with low fidelity for decisional purposes (increased internal noise). This level of understanding requires finer characterization of the mechanisms underlying STM extraction from noise.

Our goal is to gain computational understanding of how STMs are processed by normal and impaired auditory systems. To achieve this objective, we developed a novel framework based on psychophysical reverse-correlation (Ahumada & Lovell, 1991; Murray et al., 2011) that builds upon, but also goes further than, the approach adopted by Oetjen & Verhey (2015). Previous studies relying on this technique have clarified various aspects of suprathreshold auditory tasks, e.g. nonlinear mechanisms for tone-in-noise detection (Joosten et al., 2012), retuning of AM-processing (Joosten et al., 2016), modulation primitives supporting speech-in-noise intelligibility (Venezia et al., 2016, 2019). Successful application of reverse-correlation relies, among others, on two inter-related factors: (i) the nature and structure of the perturbing noise source must efficiently interfere with the mechanisms engaged by listeners for detecting the target, and (ii) a large data mass (several thousand trials) is necessary to derive a stable, accurate image of those mechanisms.

The specific data mass required to obtain a stable perceptual filter varies with several factors, including stimulus complexity and the characteristics of the perceptual process under investigation. In previous approaches measuring perceptual filters for detecting a single tone in noise, the noisy perturbation was applied to the full time-frequency domain (e.g. Shub & Richards, 2009; Joosten et al., 2012), requiring trial counts as large as 10K to obtain an interpretable image. The inefficiency of these measurements may be attributable to the fact that STM signals are complex patterns that may not be effectively masked by a time-frequency perturbation. To address this issue, we designed a low-dimensional modulation noise that can be represented by a few components, and acts as an efficient masker for other (to-be-detected) modulations (see Figure 1, panels C and D, as well as Methods for further details).

Our approach builds upon, and is inspired by, prior work in the characterization of visual processes selective for orientation (Ringach, 1998; Neri, 2014a). More specifically, prior psychophysical work has demonstrated that perceptual analyzers can be successfully characterized as oriented sensors (Adelson & Bergen, 1991), where orientation may be defined e.g. across space-space (Neri, 2015) or space-time (Burr et al., 1986; Neri, 2014b). More relevant to the present investigation, it has also demonstrated that noisy processes spanning different dimensions of the target stimulus, such as its native space-space or its orientation subspace, can be leveraged to probe the same perceptual operation and constrain a unique computational description of that operation (Neri, 2015).

The advantage of targeting the dimension(s) along which the perceptual process operates, as opposed to the dimensions along which the hardware generates the stimulus, is that the noise structure becomes far more efficient and is therefore able to expose computational characteristics (e.g. gain control) that do not necessarily become measurable using other types of noise (Neri, 2015; Neri, 2018; see also Mangini & Biederman, 2004 for a similar discussion in high-level vision, where sinusoidal noise is found to be more efficient for probing face processing as compared to pixel noise). With this evidence in mind, we designed a one-dimensional STM noise probe spanning only a small subspace of the full time-frequency space probed by prior studies (Shub & Richards, 2009; Joosten et al., 2012). The components of this noise correspond to spectrotemporal rotations of a specific STM pattern, in a manner analogous to the orientation noise used in vision (Ringach, 1998; Neri, 2014a; Neri, 2015).

This framework was deployed to probe the mechanisms underlying STM detection simultaneously for NH and HI individuals. Particularly for the purpose of testing HI individuals, the design of efficient data-collection protocols is absolutely critical, further motivating our efforts towards identifying optimal noise probes (see above). Following detailed characterization of the auditory filters engaged by NH and NI listeners, we exploit these perceptual estimates to constrain tightly integrated data-driven modeling efforts. Our results provide evidence that NH and HI individuals rely on a similar architecture of auditory mechanisms. However, while the perceptual filters engaged by NH individuals are well tuned to span the STM region corresponding to the target signal, the filtering mechanisms available to HI individuals are suboptimally biased away towards other regions of STM space. We demonstrate that these complex nonlinear filtering strategies can be well accounted for by the classic modulation filter-bank auditory model, and that a simple loss in cochlear frequency selectivity can account for the overall shift toward temporal modulations in the HI group. Interestingly, we show that the behaviors of some individuals significantly deviate from the group-level patterns, in both NH and HI groups, highlighting the additional contribution of other mechanisms for processing supra-threshold signals that varies across listeners.

## Results

Before detailing our results, we draw attention to the fact that all listeners tested in this experiment were able to successfully perform the detection task: we individually tailored target intensity to reach a performance of *d’*∼1 (76% correct responses). In presenting our results, we pool data from all experimental sessions because we observed little learning/retuning effects over time for both groups (see complementary analysis in Supplementary Material).

### Performance metrics

Psychophysical performance is primarily controlled by 3 factors: stimulus discriminability (*d’* sensitivity), response bias and internal noise intensity. We measured all three and found no substantial difference between NH and HI groups, nor any trend of specific interest. Sensitivity was comparable (NH group: *d’* = 0.7, SD=0.2; HI group: *d’* = 1.1, SD=0.6; *p*=0.04), there was no response bias (target-unrelated preference for one interval) in either group (NH group: *c*=0.1, SD=0.2; HI group: *c*=0.0, SD=0.2; *p*=0.31), and internal noise (assessed using double-pass consistency (Burgess & Colborne, 1988; Neri, 2010a), see Methods) was similar between NH (M=1.3, SD=0.4) and HI individuals (M=1.3, SD=0.5; *p* = .96). We found no correlation between internal noise and *d’* values (NH: *p* = 0.51; HI: *p*=0.44), indicating that these two metrics reflect different aspects of perceptual processing in our task (see Fig. S4A).

### Perceptual filters derived from separate analyses of target-absent and target-present stimuli

We derived perceptual filters from all noise fields (black in Fig. 2A), as well as separately from target-absent (blue) and target-present (red) stimuli (see Methods). Because we did not observe substantial differences between filters derived from participants who were asked to detect an upward-directed target as opposed to a downward-directed target, when averaging across observers we realign data from the two conditions so that target orientation always takes a notional value of 0 (center of x axis in Fig. 2A).

**Figure 2.**
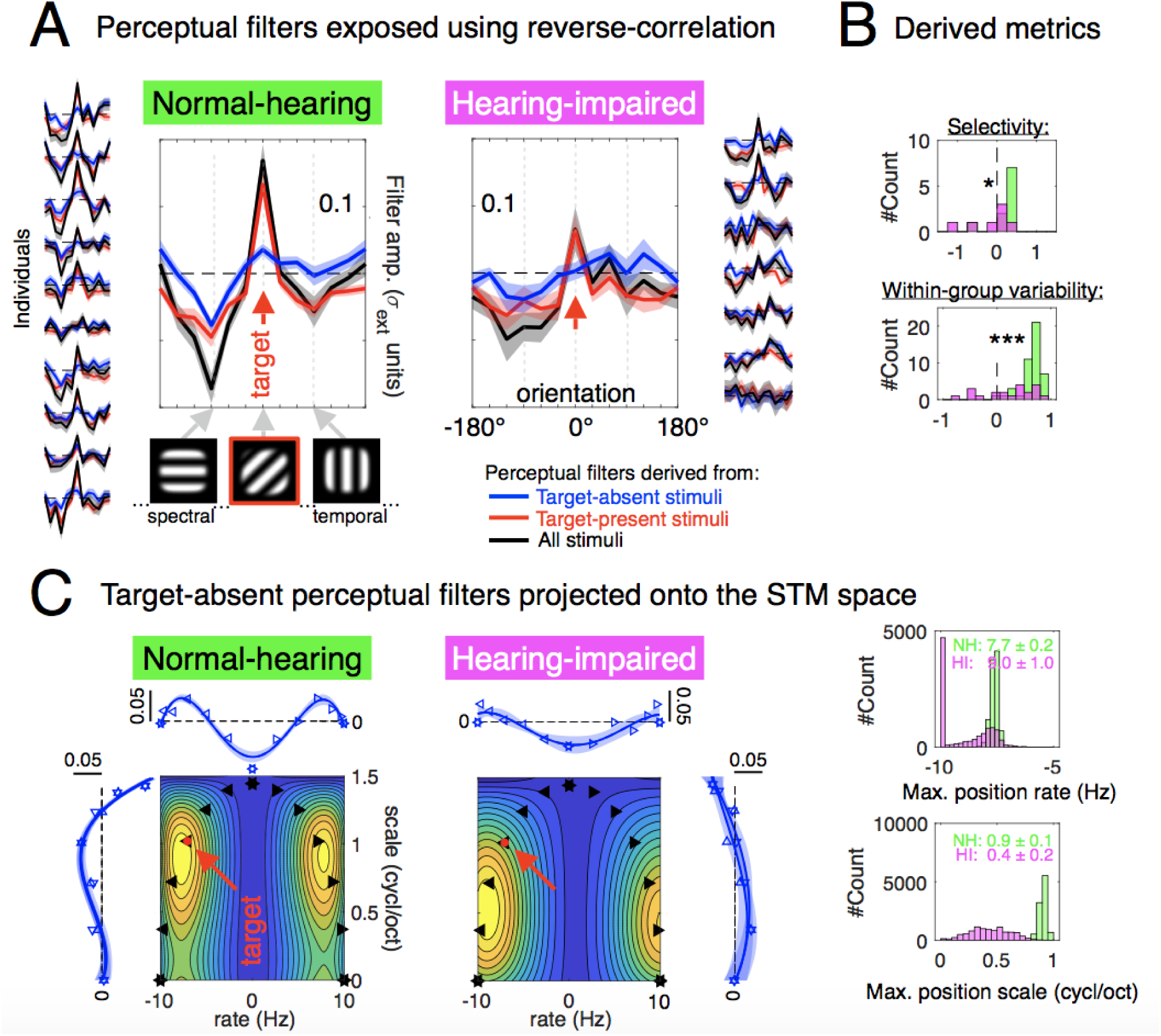
A) « Perceptual filters » derived from reverse-correlation using all trials (black traces) or data specific to target-absent or target-present stimuli (blue and red traces, respectively). Filters are plotted against the 1-D noise and centered on target orientation (red arrow). Both aggregated (middle) and individual filters (in columns) are presented for each group (NH in left-hand panels and HI in right-hand panels). Shaded areas show SEM across individuals in the middle panels or SD estimated by bootstrapping the trials for the individual traces. B) Distribution of metrics computed from the perceptual filters of NH (green) and HI (pink) individuals. The upper panel presents the introduced selectivity measure of the listeners’ perceptual filters (energy ratio at target orientation / energy at other orientations, in log-units). The lower panel shows a metric designed to capture interindividual variability within each group (correlation between each individual’s filter and the filter from every other individual of the group). Stars show significant differences between NH and HI individuals (two-sided Wilcoxon rank-sum tests; * indicate p < .05, *** indicate p < .001). C) The perceptual filters derived from target-absent stimuli are projected along the two dimensions of the native modulation space, i.e. rate and scale respectively (see horizontal and vertical traces; where shaded regions show SEM across polynomial-fitted curves on each individual). Separate projections (superimposed vertically) were derived from data points in the negative vs. positive quadrant of the MPS space (leftward vs. rightward-pointing arrows); these traces are barely distinguishable. In order to expose a more readable image of the underlying strategies of both groups, we reconstructed the full filters in the two quadrants of the MPS space by multiplying the corresponding rate and scale traces (thus assuming full separability). In each panel (left: NH; right: HI), the red arrow points toward the parameter of the target modulation (−7.1 Hz; 1 cycl/oct), which we arbitrarily projected on the left quadrant for all listeners (i.e. also for those with a downward target). While for the NH group, the filter displays band-pass characteristics well aligned with the target modulation, the filter of the HI group is shifted toward lower scale and higher rate. This shift is robust at the group-level, as shown by the distributions computed from the maximum of these patterns on the scale and rate dimensions by bootstrapped individuals (right panel).

Perceptual filters indicate how listeners differentially weight energy from different components when performing the task. A positive value indicates that more energy on this component steered listeners towards reporting that the target was present, while less energy on this component makes listeners less likely to identify the stimulus as containing the target.

We first note that, for almost every individual tested, filter estimates display significant structure (no flat patterns), clearly demonstrating that our approach is capable of exposing an image/signature of the perceptual filtering strategy adopted by listeners. The only exception is HI7 who presents with a nearly flat pattern (bottom trace in HI filters). Because this participant produced exceptionally poor performance during the first session when his threshold level was established for subsequent data collection, all data from this participant was collected for a stimulus SNR that was 6 dB higher than other HI individuals. During the following sessions, however, his performance improved to *d’*>2, a performance regime that is severely suboptimal for recovering accurate estimates from reverse correlation (Murray, 2011). Indeed, for this particular individual, the selected SNR level was too high for the external noise source to produce measurable impact on his behavior. Due to the intensive nature of data collection in these experiments, we were only able to test a limited number of individuals and are therefore not in a position to exclude data from this individual. We verified that our results remain unaffected when this individual is excluded. For similar reasons (small size of population samples), and in light of potential inter-individual differences particularly within the HI group, we present and discuss relevant effects not only at group level, but also at the individual-listener level.

On average and in both groups (black traces in Fig. 2A), perceptual filters display Mexican-hat-shaped tuning profiles: a central positive peak corresponding to the orientation of the target component, flanked by negative troughs on both sides. This shape is consistent with orientation-tuning measurements from visual tasks (Ringach, 1998; Neri, 2015). The aggregate filter from HI participants (Fig. 2A right) presents reduced amplitude compared with the corresponding measurement from NH listeners (Fig 2A left). When we consider individual patterns, we find clear deviations from the average stereotypical Mexican-hat shape, as it can be observed in the columned traces on the left and right side of Fig. 1A. In the NH group, all 10 individuals return profiles peaking at target orientation (left insets in Fig. 2A), while in the HI group some individuals produced filters peaking at orientations corresponding to higher temporal modulations and lower spectral modulations (right insets in Fig. 2A).

As a first step towards assessing listeners’ perceptual strategies in the task, we computed two overall metrics. First, we computed the root-mean-square (RMS) value of their perceptual filters. We found that this metric is negatively correlated with internal noise (non-systematic variability) in both groups, either considered separately or together (Spearman rhos, all Ps < 0.05; see Fig. S2), in line with theoretical expectations (Ahumada, 2002; Kurki et al., 2014; Murray, 2011; Richards and Zhu, 1994). Second, we computed their absolute efficiency in the task as it is typically defined: 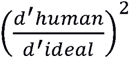 (Tanner & Birdsall, 1958; Green & Swets, 1966). Absolute efficiency values felt between <1% and 9%, which is lower to what is typically obtained in simple visual and auditory tasks (Geisler, 2003; Berg, 1990), but nevertheless consistent with valued collected in auditory tasks involving masking (Joosten & Neri; 2012; Oberfeld et al., 2013). These values did not significantly differ between both groups (NH group: *M* = 5%, SD=3%; HI group: *M* = 2%, SD=2%; *p*=0.31). Efficiency was correlated with internal noise in the NH group (*p*=0.03) but not the HI group (*p*=0.44).

Next, we constructed two different metrics (plotted in Fig. 2B) to capture the most important differences between filters from the two groups. The first metric was designed to assess the selectivity of listeners’ perceptual strategies by measuring the optimality of the perceptual filter for the assigned task (energy of perceptual weight at target orientation vs. energy of perceptual weights at other orientations, see Methods). NH individuals show greater filter selectivity than HI individuals (*p* = 0.02). The second metric was designed to assess inter-individual variability among listeners within each group. We computed the correlation between each individual’s filter and the filters from other individuals of that group, and found a significantly higher variability in the HI group as compared to the NH group (*p* < .001).

### MAX-model demonstrates non-linear processing strategy

Target-present and target-absent filters are notably different: target-present estimates present a clear modulation around target orientation that is markedly reduced in target-absent counterparts. This result is incompatible with a linear template-matching strategy (Ahumada, 1967; Neri, 2004), prompting us to adopt computational tools that can accommodate departures from the template model (Neri, 2010c).

To clarify the above statements, we demonstrate how a simple variant of the popular MAX uncertainty model (Pelli, 1985) is sufficient to reproduce the experimental estimates (see also Dau et al., 1997; Joosten & Neri, 2012). On each trial, the model weighs each of the 12 stimulus components as specified by a 12-vector template (Fig. 3A); it then takes the maximum value for each stimulus, and selects the stimulus producing the larger value. NH and HI data are modeled by different templates (green and magenta curves in Fig. 3A).

**Figure 3.**
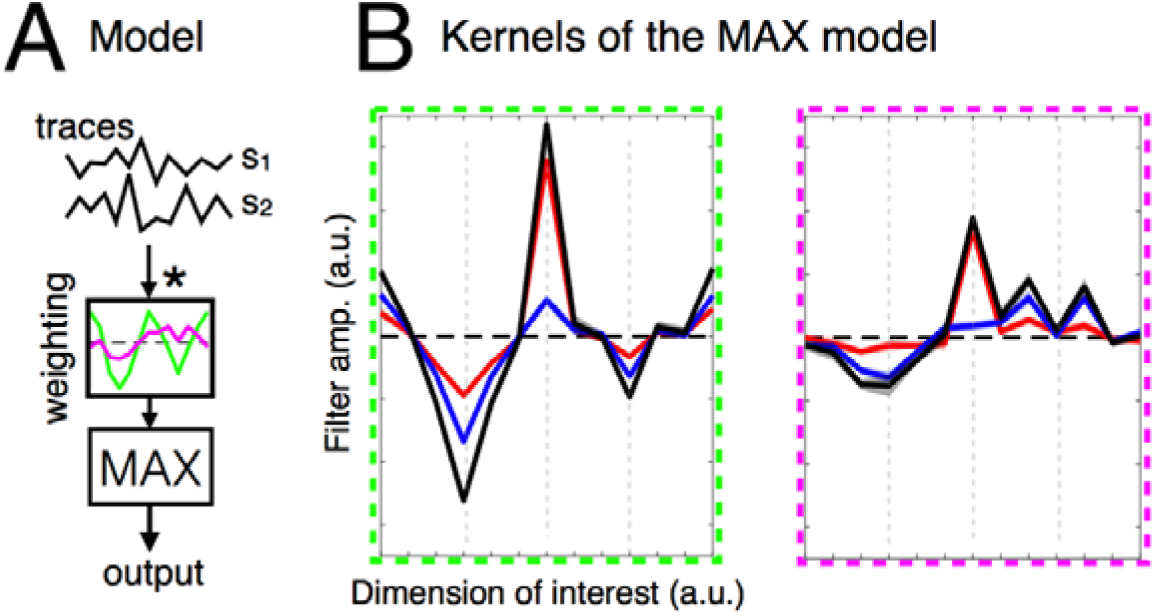
A) Structure of the MAX-model tested. On each trial, it takes as input the traces (12 components) corresponding to the energy of the different modulation components forming each stimulus s1 and s2, multiplies these traces by a given weighting profile (magenta or green curves) and picks (through a MAX operator) the stimulus that produces the highest value. B) reverse-correlation filters (same legend as in Fig. 2) obtained from this model, corresponding to the two different weighting curves proposed.

As shown in Fig. 3B, this simple model captures all main features of the data. While the choice of templates is arbitrary, the simulated difference between target-absent and target-present filters is not a consequence of this choice, and is instead produced by the MAX nonlinearity (replacing MAX with linear summation results in no difference between target-present and target-absent estimates). These simulations demonstrate that a simple MAX structure can, in principle, account for the complex pattern exposed by our measurements. Because the perceptual strategy engaged by listeners was strongly nonlinear, we cannot transparently interpret the filter estimates in terms of “perceptual weights” applied by listeners to different stimulus components (Neri, 2018b). As expected from theory (Neri, 2004; Tjan & Nandy, 2006; Neri, 2010b), simulated target-absent profiles closely resemble the model templates preceding the non-linear stage.

### Computational strategies inferred from target-absent perceptual filters

Based on the above considerations, we focused on average target-absent filters to infer perceptual weighting profiles. In the NH group, target-absent filters present a peak at target orientation; in the HI group, the peak is shifted towards temporal modulations. In both groups, target-absent filters displayed approximate symmetry around the “pure temporal modulations” line.

To aid visualization of this result, we project target-absent perceptual filters onto rate-scale dimensions and reconstruct the filters in the two quadrants of the MPS space (see Fig. 2C), assuming rate-scale separability (Chi et al., 1999; Venezia et al., 2019). Overall, the filters in negative and positive quadrants of the MPS space are symmetric, although we can notice slightly higher values for the quadrant that contains the target (which we arbitrarily projected on the left side for all subjects; see the contour plots in Fig. 2C), for both groups. In the NH group, non-directional band-pass characteristics are finely tuned to the parameters of the target modulation (peak in Fig. 2C (left) falls near target location). In the HI group, the frequency characteristics are biased towards lower scale and higher rate values (peaks in Fig. 2C (right) are closer to bottom corners). The robustness of this result is confirmed by bootstrapping across individuals (see Methods), yet it presents large inter-individual variability, particularly within the HI group.

To summarize the above results, the perceptual strategy of both NH and HI groups can be modeled as largely non-directional bandpass filters followed by a non-linear rule akin to a MAX operation. The frequency characteristics of the bandpass filters match those specified by the signal-to-be-detected in the NH group, but not in the HI group. In the latter, they are shifted towards lower scale and higher rate values.

### Filters predicted from the modulation-filterbank model

The MAX model is not intended as a physiological implementation of known facts about the auditory system. We therefore complemented the preceding simulations with a biologically-inspired computational model to identify specific deficits that may underlie the observed differences between NH and HI listeners. We tested a simplified version of the popular modulation filterbank model (Dau et al., 1997). We used default values for all parameters (see Methods), removed the earliest compressive stage (unlikely to be relevant for the task) and adopted a MAX cross-correlation device as final decisional stage (Fig. 4A, see Methods for further details).

**Figure 4.**
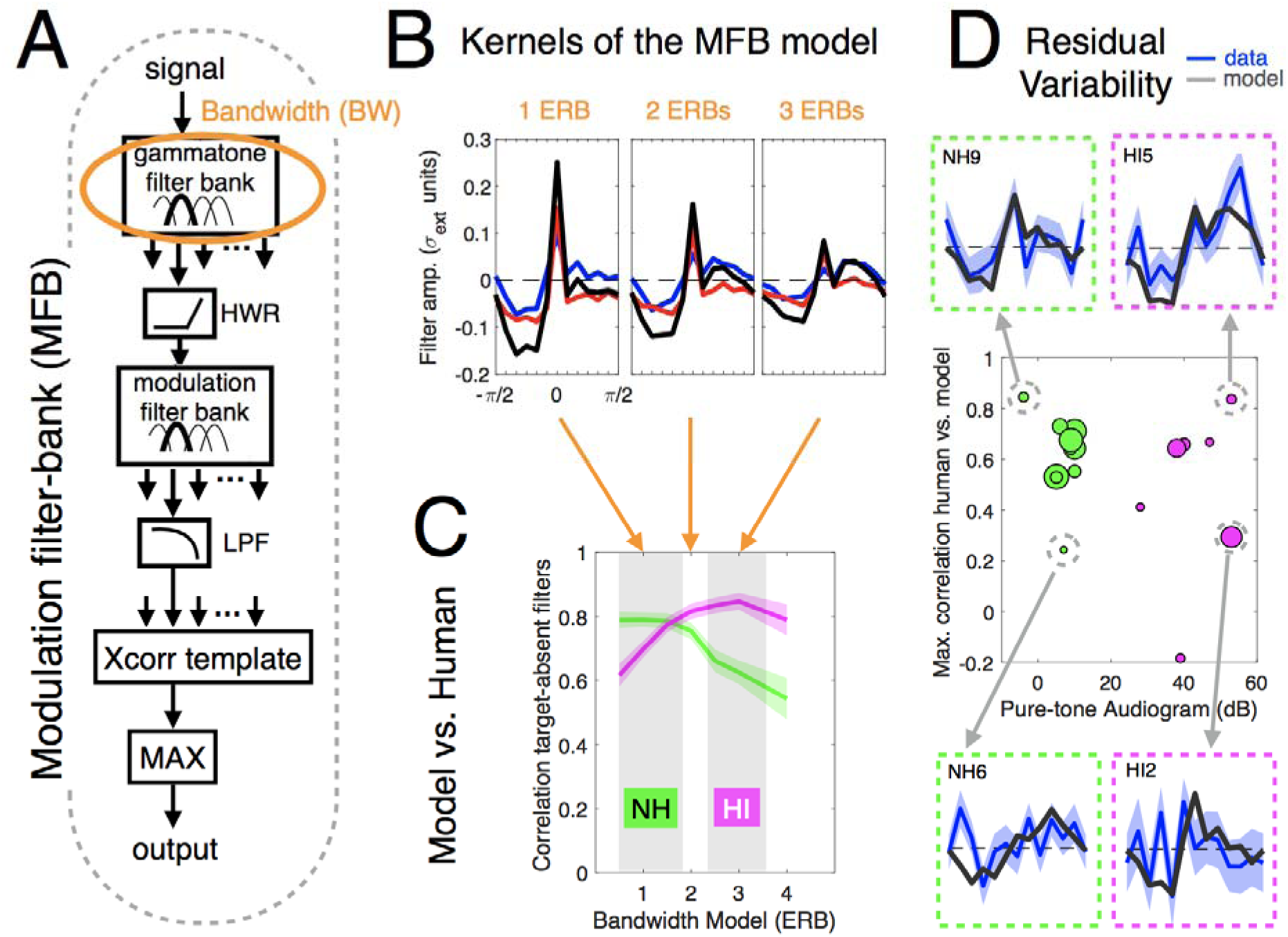
A) Illustration of the structure of the modulation filter-bank model tested (highlighted in orange: the cochlear stage). B) Examples of kernels derived from a reverse-correlation analysis of the model’s predictions using gamma-tones filters with bandwidths (BW) from 0.5 to 4 ERBs. Shaded areas correspond to SD estimated from bootstrap. C) Correlations between the model and the listeners’ target-absent filters for various BW, demonstrating a quantitatively better match for NH individuals using 0.5-1-ERB vs. 2.5-3-ERB wide BW for HI individuals. D) Maximum human-model correlations across ERBs plotted against pure-tone audiograms (averaged between 500Hz and 4kHz) shown for all NH and HI individuals (dot size proportional to absolute efficiency). This panel illustrates the interindividual variability that cannot be captured by variation of cochlear tuning in the MFB model, both for HI and NH individuals. We highlight in particular two HI individuals and two NH individuals with similar audiograms but distinct perceptual filters for STM detection: those on top are better accounted for by the model than those at bottom, a result that cannot simply be explained by differences in efficiency.

To gain some insight into how sensorineural hearing loss (SNHL) contributes to the differences observed in the HI group, we explored the effect of frequency selectivity – one particular deficit associated with SNHL (Moore, 2007; Lesica, 2018) – on the simulated perceptual filters. More specifically, we varied filter bandwidth at the cochlear stage between 0.5-ERB and 4-ERB wide (see Methods for details).

Filters derived from these simulations are presented in Fig. 4B. Qualitative inspection of these patterns shows striking similarities with those obtained from real human judgments. The simulated profiles for ∼1-ERB bandwidth (BW) reproduce the asymmetrical Mexican hat-shapes (black trace) as well as the major features of both target-present and target-absent filters (red and blue traces). In particular, target-absent filters successfully reproduce the bimodal pattern observed in human perceptual filters with a main peak at target orientation and a secondary peak at the orientation corresponding to a 90°-shift. Strikingly, simulations for 2.5-3 ERBs of BW show close resemblance with the corresponding empirical estimates of the HI group: the peak of target-absent filters is shifted towards temporal modulations, and the negative flanks on both sides appear less sharp, as observed in the average HI data.

We confirm these group-level observations by computing the correlation between simulated and measured target-absent filters at different ERB values; highest correlations (Pearson correlations ∼.8) are returned at BW = 0.5-1.5 ERB for the NH group and at BW = 2.5-3 ERBs for the HI group (see Fig. 4C). This result demonstrates that the model with normal cochlear tuning well accounts for the average pattern of the NH group, and that a two-to-threefold broadening of cochlear tuning can account for the average pattern of the HI group. Yet, we show that the present model and the variations of cochlear tuning can however not explain the behaviors observed at the individual level, for both groups (see below).

Figure 4D plots the maximum human-model correlation across all possible ERBs (ranging between .5 and 4 ERBs), i.e. the best that the model can do when allowed to vary cochlear tuning. As can be seen, there is a notable variability regarding how this model can account for individual patterns, with correlation values varying between .3 and .9. Critically, this variability is observed for both NH and HI groups, and is found to be unrelated to the average pure-tone audiograms of these individuals as well as their absolute efficiencies (all correlations non-significant, Ps > .05). To further illustrate this result, we re-plot the target-absent filters of two HI individuals and two NH individuals who exhibit distinct perceptual filters in the task, despite having similar audiograms. Critically, while the model with allowed variation of cochlear tuning can account for the behaviors of NH9 and HI5, it cannot capture as well the patterns of individuals NH6 and HI2.

## Discussion

This study capitalizes on the richness of a large dataset derived from a psychophysical reverse-correlation task specifically designed to probe the mechanisms underlying detection of auditory spectrotemporal modulations (STM) in both NH and HI listeners.

We successfully deployed a reverse-correlation approach in the STM domain by generating low-dimensional external noise that efficiently impacts listeners’ detection mechanisms. To this aim, we developed a novel framework based on 1-D STM noise created from rotations of the STM target in spectrotemporal space. All but one listener tested with this procedure reached an optimal performance regime of *d’*∼ 1 for individually tailored SNR levels, demonstrating that our protocol can be efficiently applied to both NH and NI populations.

The associated perceptual filters exhibit clear structure, further demonstrating the efficacy of our 1-D STM-noise design. These filters show different patterns for target-absent vs. target-present stimuli in both groups, exposing the presence of non-linear processes unaccounted for by a template-matching strategy (Neri, 2004; Tjan & Nandy, 2006; Neri, 2010b). These departures from template-matching are accommodated by a small cascade model consisting of a front-end STM weighting function followed by a MAX operation (see also Joosten et al., 2016). Because this cascade structure is applicable to both NH and HI groups, this result indicates that hearing-impaired listeners rely on intact circuitry for monitoring the output of their modulation channels.

### Directional symmetry (or lack thereof) in STM space

Based on the fact that target-absent filters provide a more transparent image of the filtering strategy adoped by listeners (Tjan & Nandy, 2006, Neri, 2004, 2010b), we projected these descriptors onto rate/scale dimensions and reconstructed the associated 2D filters in MPS space (assuming full separability). Our results clarify important features of spectrotemporal modulation filtering that have been only partially addressed by previous studies (see below).

First, the projection of target-absent filters onto MPS space directly supports the presence of band-pass filtering strategies for both NH and HI listeners. This finding adds to the view that the auditory system is tuned to spectrotemporal modulations (Sabin et al., 2012; Oetjen & Verhey, 2015); here we extend this result to HI individuals by demonstrating that tuning is preserved in this group. While filters are tightly tuned around target parameters for NH listeners, their center frequency is shifted toward lower rates and higher scales in the HI group.

Overall, we observe that the tuning estimates assessed in each quadrant of the MPS space are qualitatively similar, indicating that the underlying filters engaged in the task are not directional, i.e. they do not discriminate between upward and downward spectrotemporal modulations (not required by our task), consistent with behavioral masking data in humans (Chi et al., 1999) and neural responses in animal physiology (Wooley et al., 2005). Yet, a close quantitative inspection shows that the filters in the quadrant around the target (here, arbitrarily positioned on the left), possess slightly higher peaks, compared to the filters in the opposite quadrant (see Fig. 2C, in particular in the HI group). This result would be consistent with Oetjen & Verhey (2017), who found clear asymmetric masking patterns between the two modulation quadrants. They interpret their results as supporting the presence of partially selective directional filters in STM space; however the connection between masking profiles and the tuning properties of perceptual mechanisms is opaque, as demonstrated by our own data. Indeed, if we consider tuning profiles without distinction between target-present and target-absent estimates, their marked asymmetry would suggest that STM filters are at least partially selective to modulation direction. This interpretation, however, overlooks the fact that target-present estimates are distorted by the nonlinear operator. Although it is unclear whether similar nonlinear mechanisms operate within a masking design and whether their contribution may be comparable to what we observe in our data, this possibility must be given careful consideration, weakening the evidence for directional tuning supplied by masking experiments (see above).

At this stage, our study reveals that when STM processing is probed in a detection task, both NH and HI groups demonstrate limited evidence for directional selectivity, suggesting that hearing loss might not impact this aspect of spectrotemporal modulation processing. This observed symmetry is consistent with the view of separable processes across spectral and temporal dimensions, and has therefore implications for auditory modeling (Dau et al., 1997; Schädler et al., 2012). Particular attention should be devoted to this aspect by future studies.

### Potential role of internal variability

We report similar values for internal noise at ∼1.3 units of external noise in both groups, in close agreement with the estimate returned by a meta-analysis of several visual and auditory tasks (Neri, 2010). Taken together, these results confirm that HI and NH listeners differ in the STM detection task because of systematic, non-stochastic differences in their STM filtering process. These conclusions appear at odds with a recent study (Venezia et al., 2019) reporting greater internal noise in HI listeners, however the measure of internal variability adopted by that study is indirect and it is unclear how it relates to our double-pass estimates. Further experiments will be necessary to settle this issue conclusively.

### Changes in cochlear bandwidth capture group-effects

We complement our cascade-modeling approach with the adoption of a landmark auditory model, the modulation filter-bank (MFB) model, to further understand what may have caused the observed differences between NH and HI listeners. Importantly, we find that filters returned by the MFB model with default parameterization are in excellent agreement with those derived from NH participants, and that an increase in cochlear bandwidth captures the shift of band-pass characteristics observed in the HI group (Figure 4). First, these results indicate that STM processing relies primarily on frequency selectivity and temporal-envelope-based mechanisms in the modulation domain. Second, they suggest that the differences between NH and HI listeners are not trivial and may be accounted for by one component of SNHL: cochlear frequency selectivity.

We emphasize that our results must be interpreted conservatively, for the following reasons. First, they do not imply that acoustical temporal fine-structure (TFS) information was not used. While the MFB model only carries envelope information, this characteristic partially retains TFS information via FM-to-AM conversion. Additionally, the fact that our stimuli cover a wide frequency range does allow us to evaluate the potential contribution of neural TFS in the processing, which would be mostly engaged only up to ∼1 kHz (Moore, 2007). We show that a temporal-envelope based model, without any neural TFS processing stage, can account for the present data, which is consistent with the view that listeners can indeed rely only on envelope information to perform the task. Second, the MFB model does not include complex cognitive/decisional stages, making it impossible to assess the potential contribution of said factors to our results. Our finding that a simple increase of cochlear bandwidth reproduces the main characteristics of HI filters strongly suggests (but does not demonstrate) that the shift in the peak of the latent internal weighting profile is a direct consequence of the degraded cochlear frequency representation. Yet, at this stage, we cannot affirm that frequency selectivity is the unique potential source of differences in processing between NH and HI individuals.

Interpretation of our results is further hampered by the lack of age matching between NH and HI groups (individuals in the HI group were older). While group differences are most likely due to hearing loss rather than age (previous studies have found that differences in AM processing originate from HL not age; e.g. Wallaert et al., 2017), these two factors remain confounded and their contributions cannot be disentangled without additional data. Further experiments will be necessary to pinpoint the exact source of impairment for HI listeners; our study offers a fully-fledged approach to guide such efforts.

### The effect of cochlear bandwidth does not account for individual patterns

In reaching our primary conclusions, we have intentionally averaged estimates across listeners to extrapolate beyond individual idiosyncrasies and reveal common aspects of the perceptual process. However, perceptual filters present with substantial variation across individuals within each group, and these differences do not merely reflect measurement noise (see error-bars for individual traces in Fig. 1A) but instead provide meaningful information regarding the specific processes engaged by each listener. We find larger inter-individual differences for the HI group, with a wide variety of behaviors observed among individuals with similar audiometric losses, but there are also notable differences for estimates from different NH individuals.

Using model simulations, we demonstrate that the loss in cochlear frequency selectivity instantiated by the MFB model cannot account for these intra-group differences. Varying the cochlear tuning parameter results in a gradual shift of the band-pass characteristics toward pure temporal modulations (see Figure 4B), a change that is not sufficient to capture the diversity of tuning profiles observed across HI individuals (see Figure 4C). Similarly, this model is unable to account for the differences observed within the NH group. This result highlights the need to consider additional mechanisms if we want to go beyond group-level considerations of hearing deficits. We argue that the variability that remains unaccounted for by this model is informative and could actually be used to guide further efforts to pinpoint other peripheral and central components underlying supra-threshold hearing distortions in each individual. To this end, a fitting procedure involving other model mechanisms and parameters would be necessary, and would require additional data to computationally constrain their identification.

Our results show that this inter-individual variability is not related to audibility differences: we observe that, while some HI individuals show tuning characteristics that are nearly normal (i.e. similar to those obtained from the NH group) despite substantially different hearing thresholds, other HI individuals present with different filter estimates despite having similar audiograms (see Fig. 4D). These observations support the view that our filter estimates reflect aspects of auditory processing that go beyond audibility, and that the inter-individual differences observed within the HI population reflect differential engagement of suprathreshold processes.

Among such suprathreshold processes, an interesting candidate for explaining our results relates to high-level central or cognitive idiosyncratic strategies developed by HI listeners to cope with the permanent loss of fidelity associated with their internal auditory representation. However, several other supra-threshold factors are likely involved. At this stage, it is not possible to establish whether a common modeling structure such as the MFB would carry enough flexibility to capture all different filters observed across and within groups in our study. Further work with larger samples and additional tasks will be necessary to determine which model stages could account for these inter-individual differences and thus clarify their origins. The combined experimental-modeling approach introduced in the present paper offers a principled way to assess supra-threshold auditory processing in each individual based on a detailed characterization of spectrotemporal modulation encoding. Therefore, it opens new critical avenues to explore the origins of SIN understanding differences between individuals with similar audiograms.

## Materials and Methods

### Participants

We tested 10 normal-hearing (NH) participants (age range 21-37 y.o.; M=27, SD=5) with audiometric thresholds <= 25 dB HL in the 250-8kHz range in both ears, and 7 hearing-impaired (HI) participants with mild to moderate symmetrical sensorineural hearing loss (age range 59-67 y.o.; M=63, SD=3). Their audiograms, specific characteristics, and corresponding experimental conditions are displayed in Fig. S1 and Table S1. All subjects were naïve to the goals of the study. They gave their informed written consent prior to the experiment in compliance with the Declaration of Helsinki and were paid for their participation.

### Stimuli

Ripple noise was constructed by summing 12 elementary ripples of different rates/scales with different energy/phases (see Fig. 1B and 1C). Rate/scale values were selected so that the envelopes of the 12 ripple components corresponded to rotations of 15° around a target signal with rate 7.1Hz and scale 1 cycle/octave. In our plots, the target orientation is assigned a value of 0 (see Fig. 1B, middle panel). If we denote the envelope of each component at full modulation depth with notation **M**_j_ where index j ranges between 1 and 12, j=7 indicates the component corresponding to the target. These spectral and temporal modulations fall within the region of the modulation power spectrum (MPS) that is relevant for speech intelligibility (Elliott & Theunissen, 2009; Venezia et al., 2016) and are comparable to those adopted by previous studies (Oetjen & Verhey, 2015). For each stimulus, the level of each component is denoted by k_j_ and was randomly drawn from a normal distribution (SD = 3 dB, restricted to ±3 SD), while the phase of each component was assigned a pseudo-random value chosen from [0, pi/4, pi/2, or 3pi/4] (data were then pooled across the different phases, because we found no differences in the filter estimates when computed separately for the 4 possible phase values). The 12 envelopes were superimposed to generate a composite *noise* envelope **N** = ∑_j_ k_j_**M**_j_ (an example is shown in Fig. 1B, right panel). We similarly constructed a *signal* envelope **T** by setting all k values to 0 except for k_7_=ρ (amplitude of target component). The noise-only envelope **N** (target-absent stimulus), the signal-only envelope **T** (reminder stimulus) or the signal+noise envelope **T**+**N** (target-present stimulus) was smoothly tapered around the edges by a rounded-square mask to occupy a time-frequency region of 250 ms / 600 Hz-3400 Hz, and it was then applied to pink-like noise carriers made of 400 log-spaced sinusoidal frequency components with random phases spanning the 250-8000 Hz frequency region. A new carrier was generated for every stimulus. If we denote the smoothing window with **S** and a given carrier sample with **C**, this procedure simply amounts to (**T+N**)x**S**x**C** (for the target-present stimulus) where x is element-by-element multiplication (examples are shown in Fig. 1B and 1C). We emphasize that, in the expression just detailed, the signal is added at the level of the modulation envelope *before* applying the carrier. Each sample of ripple noise is represented by the 12-component vector (k_1_,k_2_,..,k_12_) which we refer to with notation **n**_i_^[q,z]^: the vector sample presented on trial i in the target-absent (q=0) or target-present interval (q=1) and that was classified incorrectly (z=0) or correctly (z=1) by the listener. For example, **n**_9_^[1,0]^ is the noise sample that was added to the target signal on the 9^th^ trial, to which the observer responded incorrectly.

### Procedure

We used a 2IFC design: on each trial, listeners were presented with both target-absent and target-present stimuli in temporal succession (but randomly ordered) and were asked to indicate which interval contained the target-present stimulus. Stimulus duration was 250 ms; inter-stimulus interval (ISI) was 350 ms. A different sample of ripple noise (k values above) was applied to the two intervals and on every new trial. The offset value applied to the target component (ρ above) was adjusted on a listener-by-listener basis through preliminary experiments measuring the value associated with stable performance of *d’* ∼ 1 (Murray, 2011). It was then kept constant for each listener throughout the rest of the experiment. The direction of the target signal (upward or downward) was randomly varied between subjects (see Table S1 for details) to verify that our conclusions remain unaffected by target direction. All stimuli were level-normalized and presented at 75 dB sound pressure level (SPL). Physical level was therefore identical between NH and HI individuals, and ensured that all frequency components were audible to HI individuals.

Sounds were generated at a sampling rate of 44.1 kHz and converted via a 16-bit resolution Meridian Explorer2 sound card. They were presented monaurally in participants’ best ear through headphones (Beyerdynamic DT 770 pro 250 ohms). Sound level was calibrated using a Bruel & Kjaer artificial ear (Type 4153, IEC318). Participants were tested individually using the exact same material (laptop, soundcard, headphones) but at two different sites: NH individuals were tested inside a double-walled sound-insulated booth in the laboratory (in Paris, FR); HI individuals were tested in a clinical environment equipped with laboratory facilities (in Reims, FR). HI individuals were not tested inside sound-insulated booths, however the average environmental noise level was low and other individuals were not allowed into the testing room during the experiments.

The experiment was divided into 6 sessions. In the first session, auditory thresholds (125-8 kHz) were measured in quiet for both ears using a Bekesy tracking procedure. Instructions were then given to the participants who were familiarized with the STM detection task. Task difficulty was progressively increased by reducing the target offset level (ρ) while monitoring performance over training blocks of 100 trials, until sensitivity decreased to about *d’*∼1 and remained stable. The associated target offset was then kept constant for the following five following.

Responses were entered via keyboard and participants received audio-visual feedback after each trial (green text + two-tone consonant chord for correct vs red text + two-tone dissonant chord for incorrect responses). Each of these five sessions comprised a first training block of 25 trials (not used for analysis) followed by 11 blocks of 100 trials. To aid participants in maintaining a stable memory representation of the target signal and to sustain their attentional level, we presented 4 repetitions of a signal-only stimulus (envelope **T** detailed above) every 25 trials. All stimuli presented across the 11 blocks were different, except for one block (randomly chosen) that was repeated twice (at a random position in the session) in order to evaluate the percentage of agreement between the two passes for the purpose of computing internal noise intensity (Green, 1964; Burgess and Colborne, 1988; Neri, 2010a).

NH listeners completed each session in approximately 60–85 minutes; sessions were separated by a minimum of 5 hours. The schedule of the experiment was slightly different for HI listeners due to time constraints at the clinic. Depending on the participant, there were between 4 and 5 slots of 90-120 min of data collection (participants were allowed as many pauses as they wished) scheduled on different days, where they could start / stop at any time during a given session and start from where they left during the following session. A total of about ∼5k trials were collected for each participant in the main task (though it slightly differed among individuals – see Table S1 for details).

### Assessment of internal noise

According to the signal-detection theory (SDT) framework, subjects’ ability to detect the target STM in the presence of external noise, here composed of other STMs, is limited by two factors: i) the systematic filtering properties of the system to filter out the different components of the external noise in order to extract the target signal (assessed using reverse-correlation, see below), and ii) the non-systematic, random variations of the system, called internal noise, which is uncoupled from external noise properties. We presented the exact same blocks of trials twice to the participants in the purpose of quantifying internal noise using the standard double-pass technique (Burgess and Colborne, 1988; Neri, 2010a). We used a signal detection theory model without bias.

Internal noise values, expressed in units of external noise SD (σ_ext_), were derived from percentage of agreement (i.e. similar responses) and percentage of correct responses measured across two passes. On average, about 500 trials were repeated twice with each individual and could be used for this analysis (see the exact number of trials obtained for each individual in Table S1).

### Reverse-correlation analysis

We used reverse-correlation to derive “;perceptual filters” engaged by listeners in our task (Murray, 2011). Filters were derived from each individual separately for “target-absent” and “target-present” stimuli to assess the presence of potential nonlinear processes: if listeners behave like template matchers, target-absent and target-present perceptual filters must match; if they differ, this result indicates the presence of nonlinear strategies that cannot be subsumed under the template-matching operation (Joosten et al. 2016). Using the notation introduced earlier and in keeping with current literature, the target-absent perceptual filter was computed as **p**^[0]^=avg(**n**_i_^[0,0]^)-avg(**n**_i_^[0,0]^) while the target-present filter was computed as **p**^[1]^=avg(**n**_i_^[1,1]^)-avg(**n**_i_^[1,0]^), where avg() indicates averaging across all trials of the indexed type. The full kernel is simply **p**^[0]^=**p**^[1]^+**p**^[0]^ and returns an image of the perceptual mechanism engaged by listeners (the link between this image and the mechanism being often opaque). All **p** estimates are normalized by σ_ext_ the standard deviation of the external noise source (3 dB).

### Statistical analyses

We used non-parametric statistics to compare the distributions of indexes derived from our measurements either against 0 or between two samples (Wilcoxon signed-rank and rank-sum tests), and explore potential correlations (Spearman rho). Due to the limited number of subjects within each group, we also used bootstrap methods to assess the robustness of our observations at the group-level, which were first inferred from the averaged data (bootstrap was conducted to build e.g. 10,000 new samples of n *subjects* from the initial pool of n *subjects*, so we could compute the indexes from the data of these new samples).

### Filters derived from the modulation-filterbank model

We compared the “perceptual filters” derived from human data with those simulated from a simplified version of the temporal modulation-filterbank model (Dau et al., 1997). The “perceptual filters” of the model were derived from its binary responses using reverse-correlation in the same way as described above for humans listeners. We used the model with the default values of the parameters (modulation filters with a Q value of 1, consistent with experimental data showing that this parameter is similar in NH and HI listeners, Sek et al., 2015; and a phase cut-off at 5 Hz for the modulation filters, Dau, 1996; Sheft & Yost, 2007), removed the earliest compressive stage, and used a MAX cross-correlation device as final decisional stage (in light of our empirical analyses, see below). To explore the effect of frequency selectivity on the estimated perceptual filters, we varied the filter’s bandwidth at the cochlear stage of the model. We estimated the “perceptual filters” of the model for the two target directions (either upward or downward) and for different cochlear bandwidths ranging from 0.5-ERB wide to 4-ERBs wide. Each combination of target direction and cochlear bandwidth was tested using a different set of 20,000 trials constructed exactly as the ones presented to the participants in the task. The SNR of these stimuli (i.e. level of offset at target orientation subtracted by the mean level of the components at other orientations, divided by the external noise SD) was similar to the one used on average with NH participants (3.3 dB), because the simulated sensitivity was within human range (*d’*∼1). The decision stage of the model consists in a template matching with an ideal representation of the target. We constructed this template by subtracting the internal representation of a ripple stimulus containing only the target component either upward (to derive the “perceptual filters” for the upward target) or downward (to derive the “perceptual filters” for the downward target) from the internal representation of a noise consisting of the same carrier without imposed modulations. This is the standard approach to construct an exemplar that will primarily reflects the modulations of the target component but not the intrinsic modulations related to the fine-structure of the carrier. A new internal representation was generated every block of 50 trials, assigned with the corresponding target direction (upward or downward) and a random phase (among the 4 possible values). In the end, the estimates obtained for the two directions and the 4 different phase values of the template were pooled together, because we did not observe systematic differences (which further corroborates our observations from human filters’ estimates).

## Acknowledgment

This work was supported by ANR-16-CE28-0016, ANR-10-LABX-0087 IEC and ANR-10-IDEX-0001-02 PSL* from Agence Nationale de la Recherche.

## Data and Code Availability

The code for stimuli construction will be made available on a public repository at the time of the publication. Data and analysis code are available from the corresponding author upon reasonable request.

## Supplementary Material

**Figure S1.**
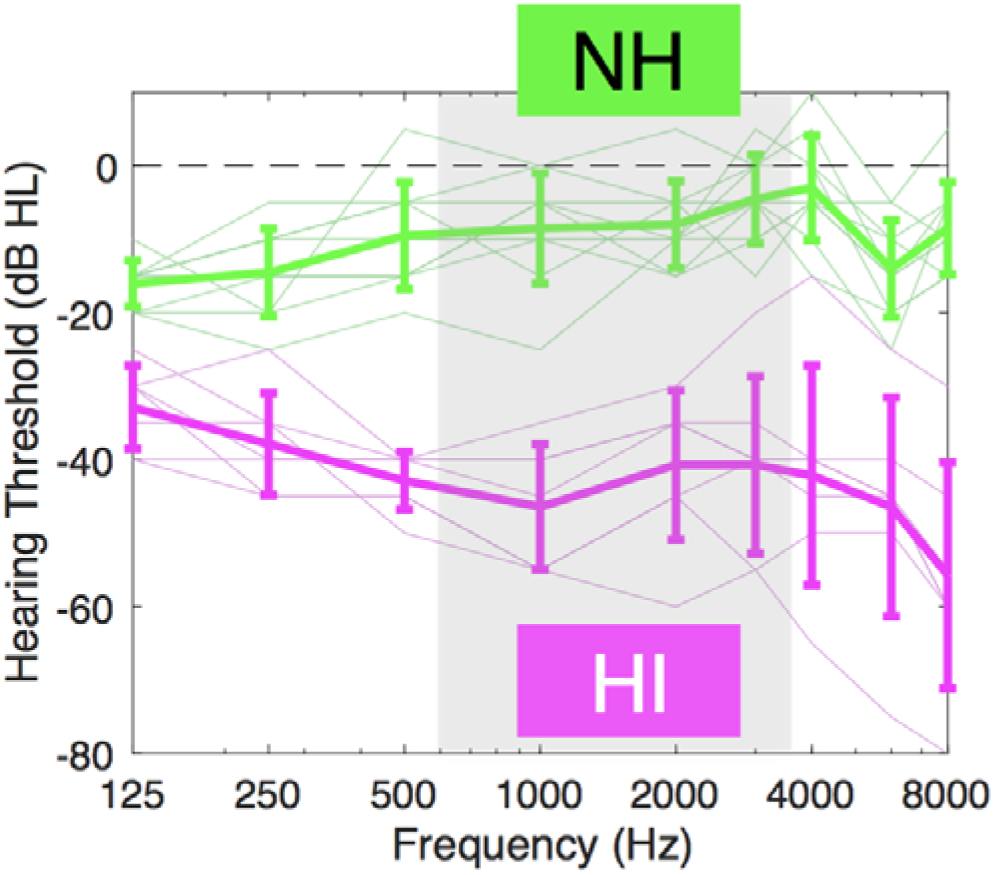
Pure-tone audiograms across frequencies in the tested ear. Thresholds were lower or equal to 25 dB HL for the participants in the normal-hearing (NH) group and around 30-60 dB HL in the hearing-impaired (HI) group. The grey shaded area indicates the frequency range of the ripples stimuli used in this study (600 Hz – 3400 Hz). Thin lines correspond to individual subjects and thick lines to group averages. Error-bars show ± 0.5 SD across listeners.

**Figure S2.**
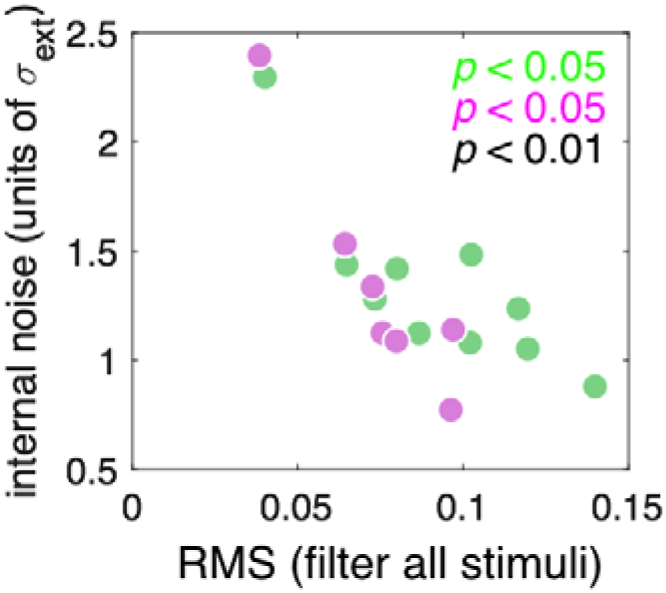
Relationship between internal noise and RMS-values of the perceptual filters derived from all stimuli in the task, for both groups of listeners (NH individuals in green, HI individuals in magenta). These values are correlated, either considering the two groups separately or together.

**Table S1.** Demographic characteristics of the normal-hearing and hearing-impaired participants of this study, and parameters of stimuli for the experiment (direction of the target, ear of presentation, SNR (dB) yielding to d’∼1, and number of trials in the task).

| participant | age | sex | target | ear | SNR (dB) | total #trials<br>(+double pass) |
| --- | --- | --- | --- | --- | --- | --- |
| NH1 | 23 | F | up | right | 2.7 | 4900 (+500) |
| NH2 | 22 | M | up | left | 2.7 | 5000 (+500) |
| NH3 | 37 | F | down | right | 2.7 | 5000 (+500) |
| NH4 | 30 | F | up | right | 3.3 | 5000 (+500) |
| NH5 | 22 | F | up | left | 4 | 5000 (+500) |
| NH6 | 28 | M | down | right | 4 | 5001 (+500) |
| NH7 | 21 | F | down | right | 4.7 | 5000 (+500) |
| NH8 | 28 | F | up | left | 3.3 | 5000 (+500) |
| NH9 | 33 | M | down | left | 3.3 | 5032 (+600) |
| NH10 | 26 | F | up | right | 3.3 | 5000 (+500) |
| HI1 | 59 | F | up | right | 4.7 | 5152 (+400) |
| HI2 | 63 | F | up | left | 6 | 4874 (+700) |
| HI3 | 61 | M | up | right | 3.3 | 4000 (+300) |
| HI4 | 67 | F | down | right | 6 | 6000 (+500) |
| HI5 | 66 | F | up | left | 5.3 | 5000 (+500) |
| HI6 | 60 | F | down | left | 12 | 5000 (+500) |
| HI7 | 63 | F | down | left | 4.7 | 3700 (+425) |

### Evaluation of potential learning and /or perceptual retuning effects

Previous studies have reported plasticity in spectro-temporal modulation processing in different populations. Based on these findings, our current study was designed so we could assess potential learning and retuning effects, which may occur and differently impact NH and HI listeners.

Indeed, Sabin et al. (2012) measured the minimal difference in modulation depth for a given spectro-temporal modulation pattern that young NH listeners could discriminate across 7 hours of task practice. They observed that their detection thresholds decreased over sessions following an exponential trend. Their study included pre- and post-tests before and after this period of task practice to evaluate how this learning would transfer to other tasks (from a discrimination to a detection task) and other stimulus conditions (i.e. STMs with other rates / scales). They found only minimal transfer effects to other conditions. Crucially, learning effects did not transfer to untrained STMs that had only one dimension in common (either spectral only or temporal only modulations), nor to STMs with a simple change in rate sign (upward to downward or vise-versa). They interpreted this result as providing further evidence that the human auditory system is composed of STM channels, i.e. joined but not separate spectral and temporal modulation channels. In other words, task practice retunes the specific channel involved to process the specific STM target concerned with the task in a more efficient way. A similar protocol deployed in old HI listeners showed learning effects of similar size but with longer time-courses and that generalized to untrained modulations (Sabin et al., 2013), suggesting a different mechanism underlying learning. Thus, we designed our study including pre- and post-tests and conducted data analyses over different experimental epochs, in order to expose specific learning or retuning mechanics that might otherwise remain obscured.

Our experiment was scheduled into six sessions. At the end of the first and the sixth session, i.e. right before and after the main reverse-correlation task, respectively, listeners participated in short pre- and post-tests specifically designed to evaluate potential learning and transfer effects (across ears and across other rate-scale values). In these tests, the stimuli were constructed similarly as in the main task (ripple noise with an offset on the target component on the target-present stimulus) and the task was identical as in other sessions. These pre- and post-tests, lasting a total of ∼25 minutes, comprised 4 different blocks made of 100 trials, which are depicted in Fig. S3: block A was identical to the main task (target ±7.1Hz, 1 cycl/oct, same ear, in block B parameters were unchanged but the stimuli were presented in the opposite ear, in block C the tested ear was the same but the sign of the rate of the target was changed, and finally in block D the tested ear was the same but the parameters of the target were modified (target: ±11.3 Hz, 1.6 cycl/oct); please refer to Fig. S3. The order of these 4 blocks was random for each subject, but the stimuli and the order in which they were presented were identical between pre- and post-tests for a given participant so as to minimize the contribution of intraindividual variability related to order effects in our measurements (Sabin et al., 2012).

**Figure S3.**
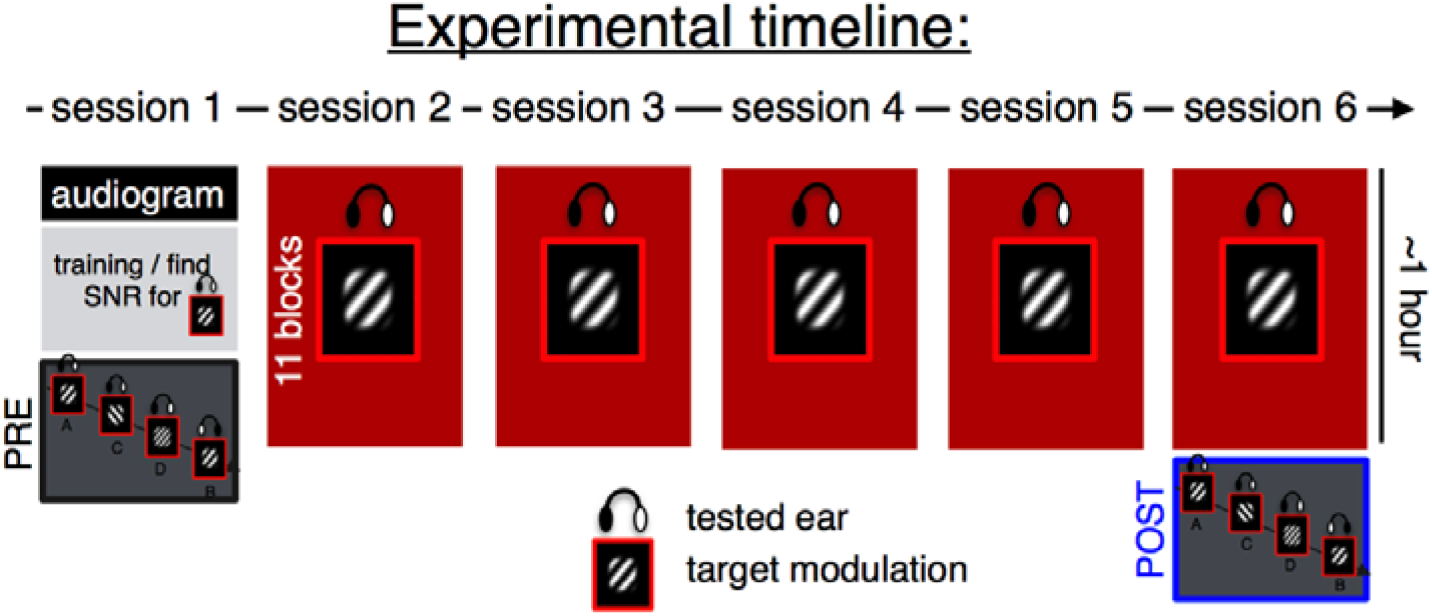
Timeline of the experiment spanning 6 sessions. The main part of the experiment consist of ∼ 5 hours of data collection (red blocks in sessions 2-6) where the same target and same ear of stimuli presentation are used – this is part for reverse-correlation analysis. The pre- and post-tests are designed to evaluate potential learning/transfer effects during the main experiment, and therefore happen right before and after it, respectively (see black and blue boxes in session 1 and 6). These tests comprise 4 different blocks (labeled A to D) to assess potential learning and transfer effects regarding the ear of presentation and direction or parameters of the target modulation. As an example, this figure illustrates the procedure for a particular subject where, in the main task, stimuli were delivered to the right ear and the target modulation to be detected was upward.

### Direct evaluation of learning effects

The performances derived from participants’ responses in the four conditions (blocks) of these tests are presented in Fig. S4. We observed better performances in the HI group compared to the NH group, and an increase in performance in post-tests compared to pre-tests in almost all conditions in both groups. A mixed rmANOVA [Group x Position (Pre/Post) x Condition] showed a main effect of Group (*F*(1,15) = 5.6, *p* = .03, □*_p_^2^* = 0.27), a main effect of Position (*F*(1,15) = 9.0, *p* = .01, □*_p_^2^* = 0.37) and a significant Group x Condition interaction (*F*(3,45) = 3.1, *p* = .04 □*_p_*^2^ = 0.17, ε = 1.00). We conducted post-hoc t-tests to compare performance in post-tests vs pre-tests for each condition. We found significant differences in condition B (t(9)=3.1426, p=0.01) and C (t(9)=3.3185, p=0.009) for the NH group, which remained significant after Bonferroni correction with n=4. There were no significant differences in the HI group (all Ps>0.05).

As in the main reverse-correlation task, we found that the performance of HI individuals was higher than the performance of NH listeners. This result can be explained by the fact that the preliminary tests to find a stable SNR corresponding to a performance of *d’* ∼ 1 were more efficiently deployed in NH compared to HI participants. However, we are interested in changes between pre- and post-tests, not in the initial differences in scores that simply reflect the success of our procedure to target the most efficient SNR value to deploy the reverse-correlation task. We observed a small increase in scores for post-compared to pre-tests, but post-hoc t-tests could statistically only confirm these differences in conditions B (opposite ear) and C (opposite target direction) in the NH group. The fact that the difference was not significant in condition A, which was the main condition in the reverse-correlation task, and that the performance improved in other conditions is not easy to explain. Our interpretation is that, although significant, these specific differences remain small and we should primarily discuss the consequences of the main effect observed: on average, all participants were performing better in post-compared to pre-tests. We believe that this benefit in post-tests simply reflects the fact that participants could better understand the task after 5 hours of practice, but does not reflect a specific learning effect. Indeed, this increased performance should have also been visible during the time-course of the reverse-correlation task, and this was not the case. Overall, these results support the view that that there was no learning during the task. If there was, it was small and likely mainly driven by task practice as we found no specific benefit to process the modulation target parameters after the 5 sessions of data collection (on the contrary the only significant differences between pre- and post-tests were found in condition B and C in NH listeners, not in condition A that corresponds to the condition of the reverse-correlation task). This means that although listeners were detecting a specific spectrotemporal modulation target embedded in noise with feedback during ∼5 hours, there was no notable improvement in their strategy to extract the target from noise. At first sight, this result might seem disappointing in terms of benefit of the task for perceptual training / learning. This seems to be at odds with studies demonstrated a specific learning to process spectrotemporal modulation (Sabin et al., 2012, 2013). However, their experimental design was different: they measured the minimal modulation depth for detecting/discriminating a given spectrotemporal modulations. Learning processes are complex: Sabin et al. (2012) observed that learning on depth discrimination for a given STM led to worsening on the detection of the same STM signal. In our experiment, listeners had to detect an offset of energy in a specific STM channel in the presence of energy in many other STM channels. This could be a first reason explaining why we did not observe any learning between the beginning and the end of the experiment. Note that previous studies using masking modulation paradigms found plasticity for AM-processing (Joosten et al., 2016) but not with STM-processing (Oetjen & Verhey, 2015). Second, it is possible that the learning occurred extremely rapidly during the initial training phase or pre-test (see Grant et al. 2013) and could therefore not be exposed here, since our pre- and post-tests were scheduled at the end of the training session and the final session, respectively.

**Figure S4.**
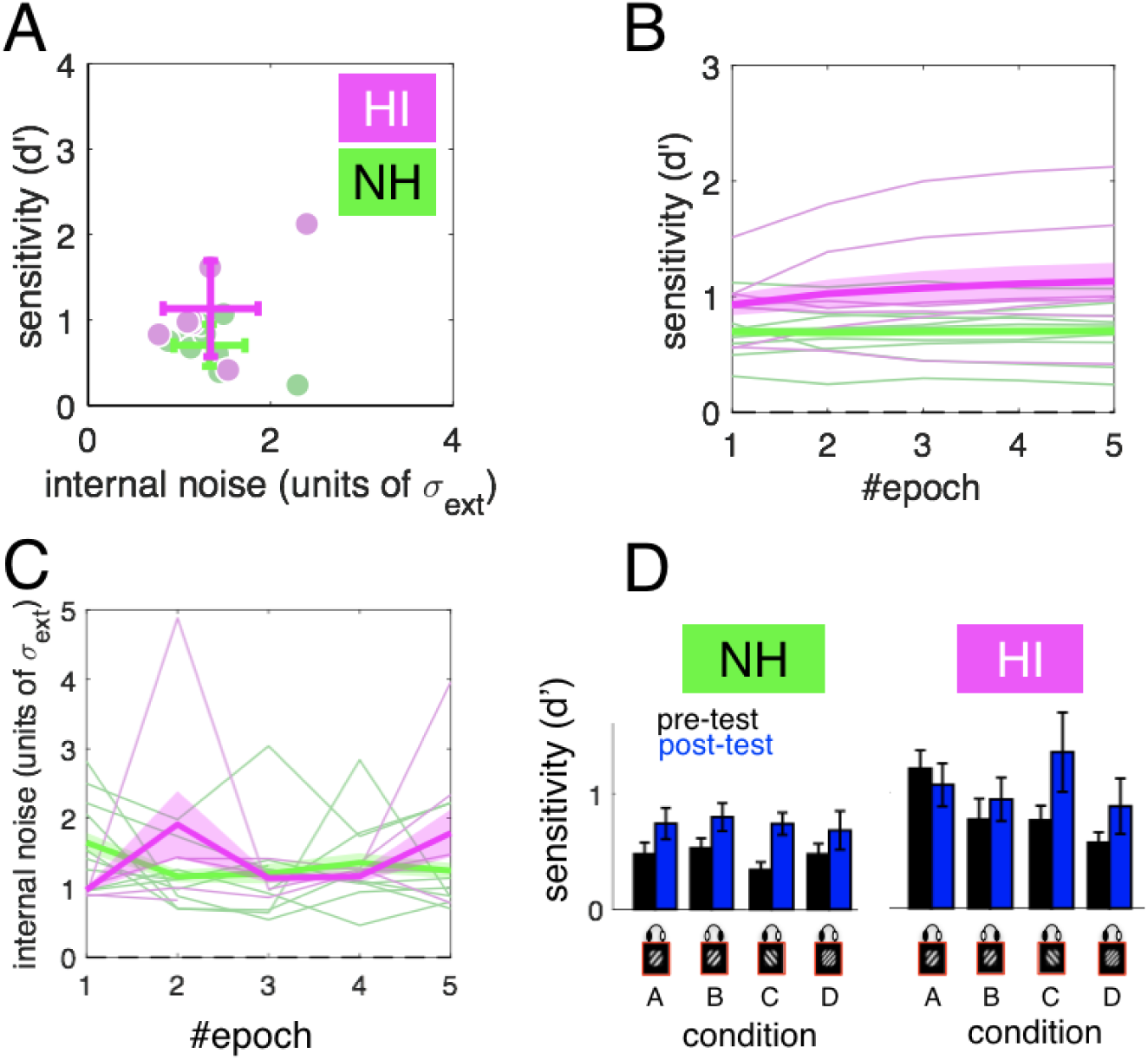
A. Values of sensitivity against internal noise for every participant of the two groups in the reverse-correlation task. Error-bars indicate mean and SD across the 2 dimensions. Panels B and C display how these two measures evolve across the experiment when data are splitted into 5 different epochs. Thick lines and shaded areas show mean and SEM in each group, thin lines show individual subjects. D. Performance derived from the 4 different conditions in pre-tests (black bars) and post-tests (blue bars) for both NH and HI participants. Error-bars show SEM.

### Evaluation of learning and/or retuning effects during the main task

In order to look for potential learning or retuning effects that would have occurred during the main reverse-correlation task, we splitted our dataset collected during the main task (sessions 2-6) into 5 different epochs and conducted the same analyses of performance, internal noise and derived the perceptual filters on the data from these 5 different epochs.

We did not observe any change in performance or internal noise across the 5 epochs in either group on average (see Figure S4), except for two HI individuals who also had the highest performance scores. The perceptual filters derived for each of these 5 epochs (not shown) were also very similar. All together, these analyses indicate that there were no systematic learning or perceptual retuning observable across the different epochs of the reverse-correlation task. This result supports the findings from analyses of pre- and post-tests that indicate no noticeable learning effects.

Then, we wished to evaluate whether the amount of data of ∼ 5000 trials per listener was sufficient to correctly approximate listeners’ perceptual filters. To do so, we computed the correlation between filters derived using subpart of data from trial 1 to N and filters derived using all trials, and varied the value of N from 250 to 5000 by steps of 125 trials. The correlation curves derived between the perceptual filters computed across restricted epochs of increasing size (from trial 1 to trial N) and the perceptual filters computed from the entire dataset are plotted in Fig. S5, in panels A, B and C. They showed a clear logarithmic growth suggesting that the number of responses collected per listener was reasonable. Yet, the fact that that a plateau was nearly but not reached also indicate that a better estimate could have been obtained with an increased number of trials. Moreover, we observed that the curve was higher for NH listeners than for HI listeners. This effect is likely due to a larger short-term variability of HI perceptual filtering strategies. Combined with the observation that there was no difference in terms of internal noise between NH and HI listeners, this result further implies that this short-term variability is due to systematic differences in processing, i.e. that the perceptual strategies were fluctuating more over time in HI compared to NH listeners.

**Figure S5.**
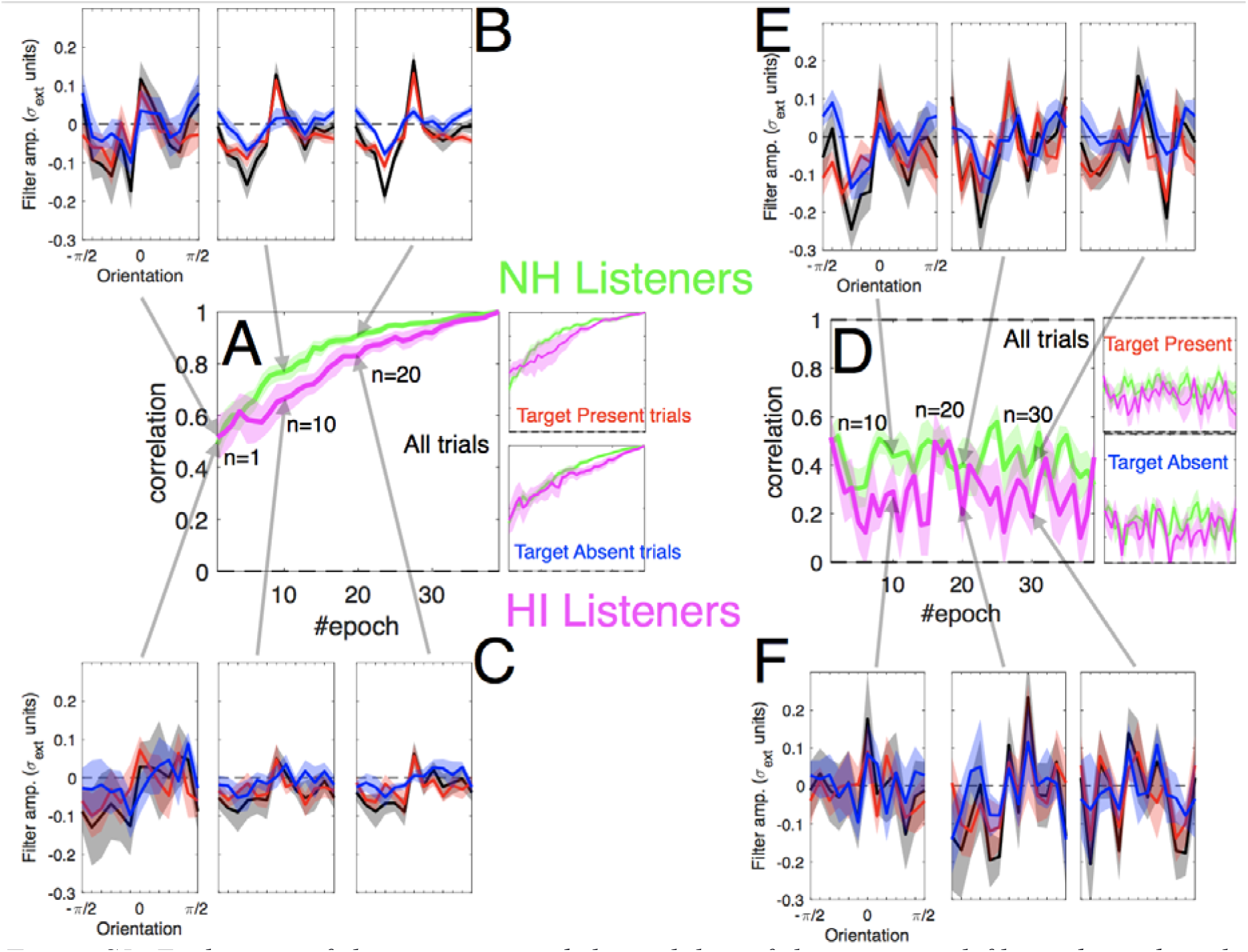
Evaluation of the precision and the stability of the perceptual filters derived in the reverse-correlation task. Panel A shows how the filters computed on a subpart of the data (evaluated onto windows of increasing size, with 50% overlap) correlate with the final filters. Panels B and C display the perceptual filters derived at specific points in both groups (top: NH; bottom: HI). Panel D shows how the filters computed on a subpart of the data (evaluated onto windows of constant size of 250 trials, with 50% overlap) correlate with the final filters. Panels E and F display the perceptual filters derived at specific epochs in both groups.

Finally, to explore for a potential retuning of the filters across time, we computed the correlation between filters derived using restricted epochs and those derived using all trials. We did not observe any trend in one direction or the other but only fluctuations, suggesting that long-term retuning or reweighting across the time-course of the experiment was absent or very small. This is shown in panels D to F in Fig. S4 where the perceptual filters were computed over short epochs along the experiment, displaying the same overall structure across different epochs.

Overall, all these results imply that an overall analysis based on the data pooled from the 5 sessions, as we did in this study, did not hide any important change or trend that would have occurred. Second, they show that our protocol does not induce measurable learning or retuning effects and cannot be considered as a potential candidate for perceptual learning experiments / to improve speech-in-noise listening strategies.

